# Impact of elevated temperature on immunity-related hormone signaling in tomato plants

**DOI:** 10.1101/2025.07.20.665745

**Authors:** Karen Liu, Vanessa Shivnauth, Christian Danve M. Castroverde

## Abstract

Molecular mechanisms governing the plant-pathogen-environment “disease triangle” are starting to emerge, although less so in agriculturally important species like tomato (*Solanum lycopersicum*). Here we analyzed defence hormone responses of tomato plants infected with the bacterial pathogen *Pseudomonas syringae* pv. *tomato* (*Pst*) DC3000 under two different temperatures. Our results showed that tomato plants exhibited temperature-sensitive expression of marker genes associated with salicylic acid (SA), jasmonic acid (JA) and abscisic acid (ABA) pathways, but not ethylene (ET). Our findings highlight the complexity of plant-microbe interactions and the importance of considering environmental conditions when studying plant defence responses.

## Description

Climate change is a serious threat to plant health, profoundly affecting the occurrence and severity of plant diseases (Altizer et al., 2013; Velásquez et al., 2018; Burdon and Zhan, 2020; Chaloner et al., 2021; Yang et al., 2022). Based on the “disease triangle” paradigm underpinning plant-pathogen-environment interactions, disease occurs through the combination of susceptible hosts, virulent pathogens and favourable environmental conditions (Colhoun, 1973; Velásquez et al., 2018). Sub-optimal environmental conditions like warming temperature can compromise plant immunity, leading to reduced resistance to pathogens (Colhoun, 1973; Zarratini et al., 2020; Son and Park, 2022; Roussin-Léveillée et al., 2024).

In response to pathogen infections, plants have developed sophisticated defence mechanisms. Plant hormones like salicylic acid (SA), jasmonic acid (JA), ethylene (ET) and abscisic acid (ABA) are important lynchpins for plant immunity (Pieterse et al., 2012; Bürger and Chory, 2019). SA is a critical hormone mediating both local basal disease resistance and systemic acquired resistance (SAR), especially against biotrophic/hemibiotrophic pathogens (Ding and Ding, 2020; Peng et al., 2021; Spoel and Dong, 2024). On the other hand, JA and ET predominantly mediate defences against necrotrophic pathogens (Glazebrook, 2005; Pieterse et al., 2012). Finally, while ABA has typically been associated with abiotic stress responses, it also plays important roles in plant pathogenesis (Lievens et al., 2017; Hu et al., 2022; Roussin- Léveillée et al., 2022).

The effects of changing temperatures on plant hormone pathways during pathogen infection have been investigated in several studies (Huot et al., 2017; Kim et al., 2017; Li et al., 2019; Kim et al., 2022; Li et al., 2024; Shields et al., 2025). However, these studies have generally focused on the model dicot species *Arabidopsis thaliana*, while the temperature-mediated regulation of plant hormone biosynthesis and signaling in agriculturally important crop species remain largely unexplored. In this study, we aimed to determine the effects of warming temperatures on defence mechanisms in tomato plants (*Solanum lycopersicum*). We specifically analyzed plants infected with the model bacterial pathogen *Pseudomonas syringae* pv. *tomato* (*Pst*) DC3000 (Xin et al., 2013) at either 23°C (ambient) or 32°C (elevated temperature) and then measured expression levels of defence hormone marker genes (Singh et al., 2021).

As shown in Figure 1, relative gene expression profiles of pathogen-inoculated tomato plants were compared to mock-treated plants (negative controls). We found that the SA pathway was temperature-sensitive in *Pst* DC3000-infiltrated tomato plants. Transcript levels of the SA marker gene *SlPR1* were significantly induced after *Pst* DC30000 infiltration at 23°C, but this pathogen-induced expression was lost at 32°C (Figure 1A). Similarly, we found that the JA pathway was temperature-sensitive in tomato plants in response to *Pst* DC3000 infiltration. As shown in Figure 1B, gene expression of the JA marker gene *SlLOXD* was only induced by pathogen infection at 23°C but not at 32°C. In contrast to the SA and JA marker genes, the ET marker gene *SlETR1/2* in tomato plants did not change across temperature or infection treatments (Figure 1C). Finally, the ABA pathway in *Pst* DC3000-infiltrated tomato plants was shown to be sensitive to changing temperature. As shown in Figure 1D, ABA marker gene (*SlLE4*) expression levels were induced by *Pst* DC3000 infection only at 23°C, but this pathogen-mediated induction was absent at 32°C.

**Figure 1.**
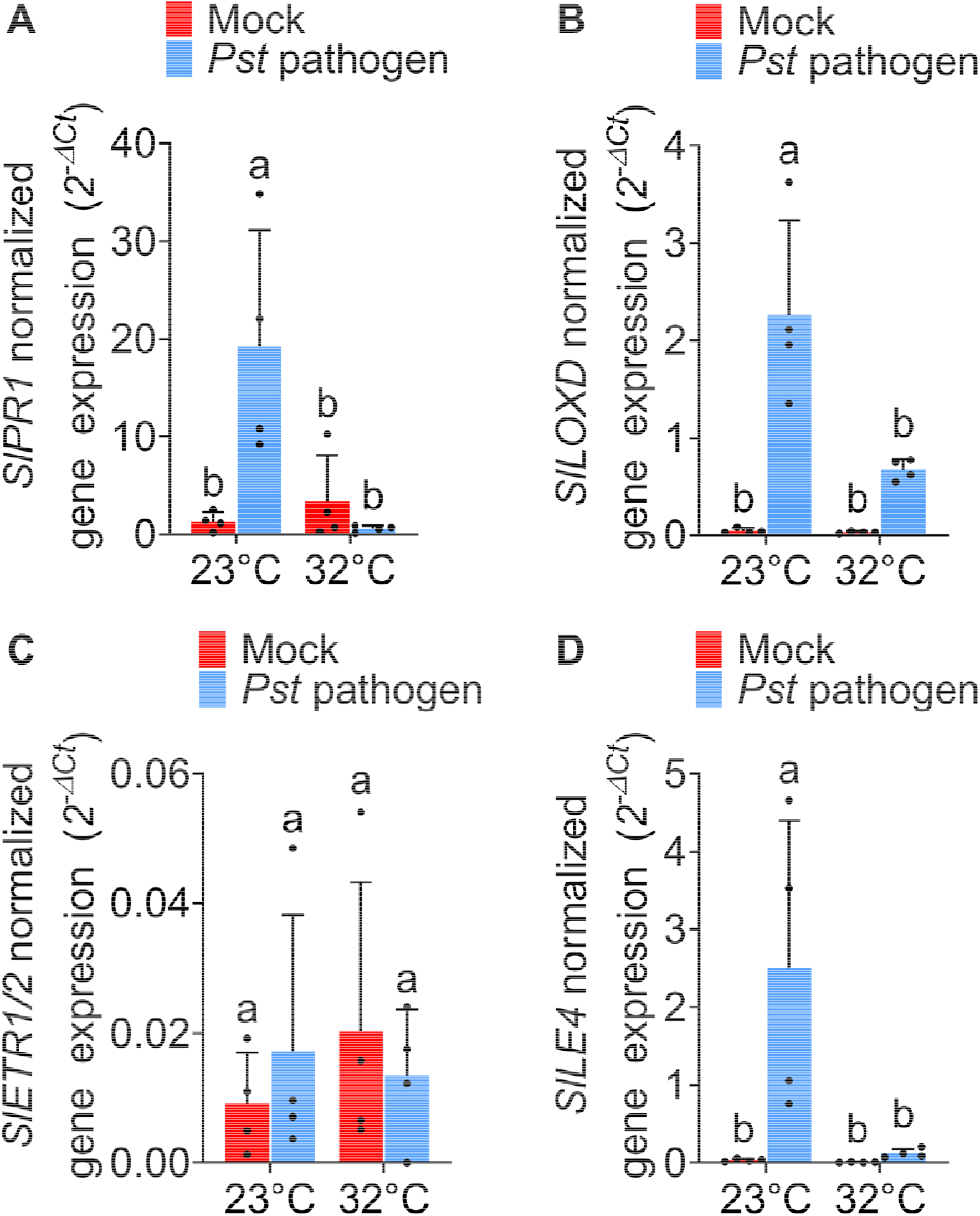
Tomato defence gene expression in response to *Pseudomonas syringae* pv. *tomato* (*Pst*) DC3000 under different temperatures. Four to five-week-old tomato plants grown at 23°C were leaf-infiltrated with either mock solution (0.25 mM MgCl_2_) or *Pst* DC3000 (OD600=0.001) and then incubated at 23°C or 32°C. Mock- or pathogen-infiltrated leaves were collected at one-day post inoculation (dpi). Leaf total RNA samples were extracted and utilized as templates for RT-qPCR analysis. The data displays the mean of gene expression values ± standard deviation for (A) *SlPR1*, (B) *SlLOXD*, (C) *SlETR1/2* and (D) *SlLE4* relative to the internal control gene *SlActin2* (n=4 individual plants). Statistical analysis was conducted using a two-way ANOVA (*p* < 0.05) with Tukey’s multiple comparisons test. Different letters demonstrate significantly different treatments. The experiment was performed 2-3 times with reproducible trends.

Temperature-sensitive induction of tomato *SlPR1* gene expression is consistent with previous studies in *Arabidopsis* and tomato plants (Mang et al., 2012; Huot et al., 2017; Kim et al., 2022; Rossi et al., 2023). However, the specific mechanisms behind the temperature-sensitivity of the tomato SA pathway remains unknown. In *Arabidopsis*, temperature-regulated GBPL3 defence-activated condensates (GDACs) govern the transcription of master immune regulatory genes *CBP60g* and *SARD1* important for SA biosynthesis (Kim et al., 2022). Given the recent discovery of tomato homologs of CBP60g and SARD1 (Shivnauth et al., 2023), as well as GBPL3 (Huang et al., 2021), a similar mechanism may be involved in tomato plants.

In contrast, our discovery of JA marker gene (*SlLOXD*) downregulation at high temperatures in *Pst* DC3000-infiltrated tomato plants contrasts with certain studies (Huot et al., 2017; Havko et al., 2020), presumably because the temperature regulation of the JA pathway may be species-specific and/or condition-specific. JA gene expression was upregulated at high temperatures in *Arabidopsis* after *Pst* DC3000 infiltration (Huot et al., 2017), in rice after *Magnaporthe oryzae* infection (Qiu et al., 2022) and in tomato after wounding (Havko et al., 2020). Nonetheless, other studies are consistent with our findings, including results showing that JA metabolism was disrupted at high temperatures in uninfected cotton (Khan et al., 2020) and *Arabidopsis* (Zhu et al., 2021). The downregulation of the ABA marker gene *SlLE4* in *Pst* DC3000-infiltrated tomato plants at elevated temperature also contrasts with a previous study in pathogen-infected *Arabidopsis*, which showed ABA marker gene upregulation (Huot et al., 2017). Finally, we found that the ET marker gene *SlETR1/2* is temperature-insensitive in *Pst* DC3000-infiltrated tomato plants. The *SlETR1/2* gene expression level was very low, which could be due to *Pst* DC3000 being a hemibiotrophic pathogen (Xin and He, 2013), while ET mediates defences against necrotrophic pathogens (Glazebrook, 2005; Bürger and Chory, 2019). However, other studies have shown temperature-mediated ET pathway changes but in different plant organs and without pathogen infection (Atta-Aly, 1992; Jegadeesan et al., 2018). The seemingly conflicting trends may be due to different factors, such as experimental conditions and plant species examined.

Although this study has shed light on the temperature regulation of plant defence hormone pathways, our results in tomato (a dicot) may not universally extend to all crops, especially major monocot plants (e.g. rice, wheat, maize). Additionally, *Pst* DC3000 was used to interrogate the effects of warming temperatures on plant defence pathways, so gene expression trends may not be applicable to all pathogens. Finally, we only focused on four hormone marker genes, so other hormone pathways could be investigated in the future. Detailed hormone quantification and global transcriptome analyses will unveil a more comprehensive portrait of defence hormone biosynthesis and signaling in tomato plants under changing temperatures.

Collectively, our findings provide a first step towards narrowing the knowledge gap on how warming temperatures affect defence hormone pathways in crop plants during bacterial pathogenesis. Together with studies in *Arabidopsis* and other model plant species (Huot et al., 2017; Cohen and Leach, 2020; Castroverde and Dina, 2021; Kim et al., 2022), this study contributes to the emerging theme of dynamic regulation of the plant hormone and immune landscape under changing environmental conditions. These mechanistic clues are critical foundations towards enhancing plant stress resilience to a warming global climate.

## Methods

### Tomato plant materials and growth conditions

Tomato (*S. lycopersicum L*.) cultivar Castlemart seeds were grown based on a previously published procedure (Shivnauth et al., 2023). Briefly, seeds were sterilized in 10% bleach at room temperature (21-23°C) for 15 mins and then rinsed 5X with autoclaved water. A final 10 mL of autoclaved water was added to the seeds, which were left to imbibe at room temperature (21-23°C) overnight. Imbibed seeds were allowed to germinate in the dark for 5 days on a sterile 9-cm Whatman filter paper inside a petri dish. Successfully germinated seeds were sown individually in autoclaved soil (3 parts Promix PGX and 1 part Turface) contained in pots (9.7cm x 9.7cm). Tomato seedlings were initially fertilized with 100mL of MiracleGro (4 g per 1 L of water) and then grown in environmentally controlled chambers (23°C, 60% relative humidity and 12h light/12h dark photoperiod with 100 ± 20 umol m^-2^ s^-1^ PPFD). Plants were fertilized weekly with Hoagland’s solution and watered regularly.

### *Pst* DC3000 pathogen infection

Four-week-old tomato plants were leaf-infiltrated using a needleless syringe with the mock treatment (0.25 mM MgCl_2_) or pathogen treatment of the model bacterial pathogen *Pst* DC3000 in 0.25mM of MgCl_2_ (OD600= 0.001) (Xin et al., 2013). *Pst* DC3000 was previously cultured in modified LB media, based on previous studies (Huot et al., 2017; Kim et al., 2022). Leaf-infiltrated plants were then grown at normal (23°C day/23°C night) or elevated temperature (32°C day/32°C) with the same relative humidity and light intensity conditions stated above. Four individual plants were used as independent biological replicates per treatment.

### Gene expression analyses

Mock- and pathogen-infiltrated leaves were harvested 24 hours after treatment. Expression levels of hormone signaling genes were quantified based on previously published protocols (Shivnauth et al., 2023; Rossi et al., 2024). Tomato leaves were homogenized with a Qiagen TissueLyser II (25 beats/s for 1 minute), with total RNA extracted using the RNeasy Plant Mini Kit (Qiagen). After measuring RNA yield and quality using a DeNovix Nanospec, RNA samples were uniformly diluted and used as templates for cDNA synthesis with the qScript cDNA super mix (Quantabio). The synthesized cDNAs were mixed with PowerTrack SYBR Green master mix (Life Technologies), and quantitative polymerase chain reaction (qPCR) was performed on the Applied Biosystems QuantStudio3 platform (Life Technologies). qPCR analyses were carried out with three technical replicates for each biological sample. Cycle threshold (Ct) values were obtained for the genes of interest and *SlActin2* housekeeping reference gene. Transcript levels of the target genes were reported as 2^−ΔCt^, where ΔCt is Ct _targetgene_–Ct_*SlActin2*_. The qPCR primer sequences are shown below.

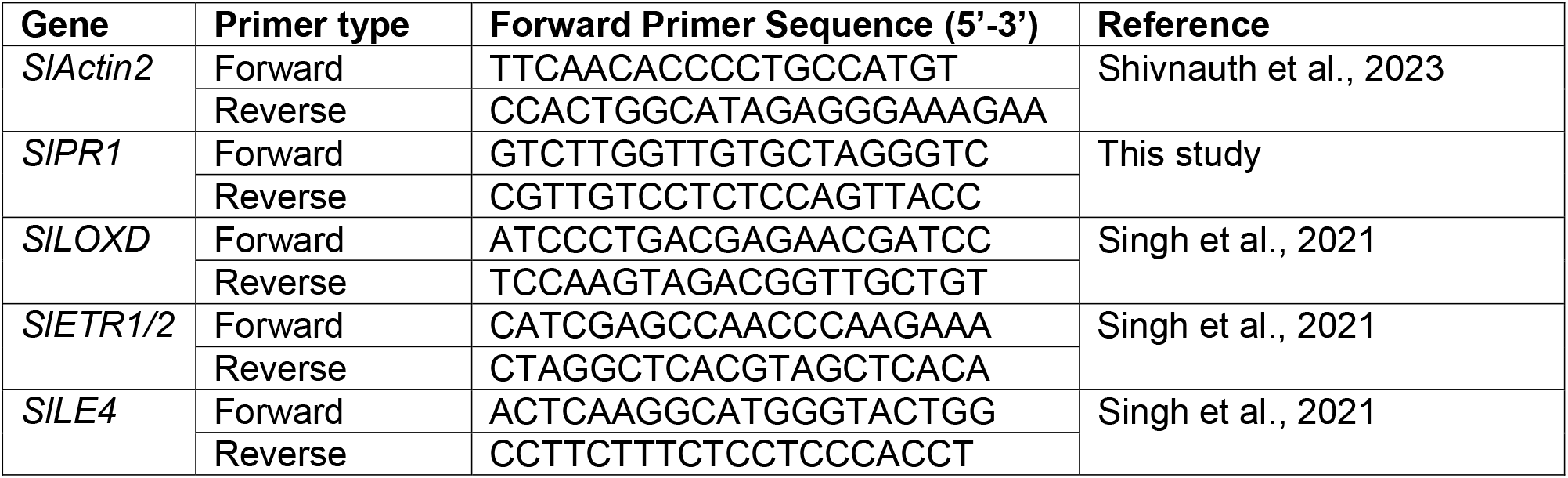

## Reagents

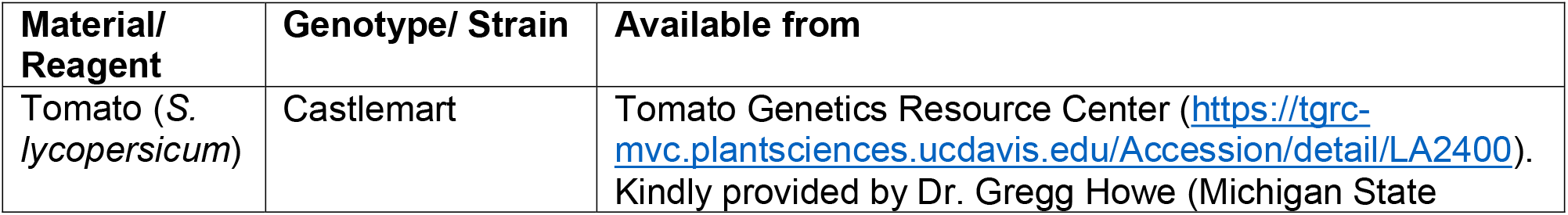
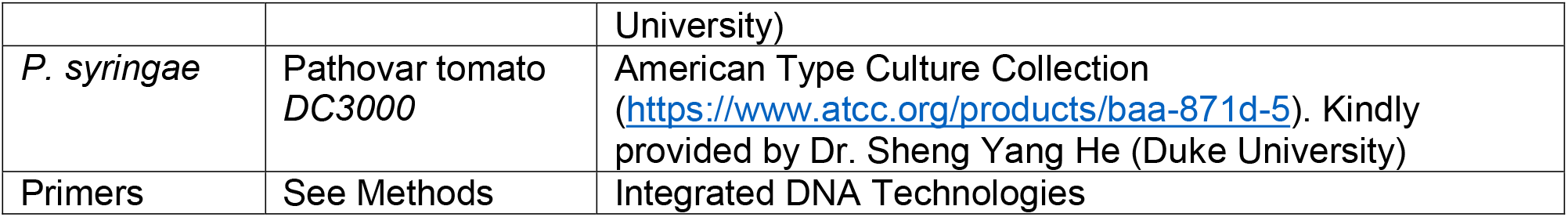

## Funding and Acknowledgements

We would like to thank Eric Marchetta and Alyssa Shields for technical assistance. Funding for this project was provided by the NSERC Discovery Grant, Canada Foundation for Innovation, Ontario Research Fund and institutional research start-up funds. Vanessa Shivnauth was partially supported by a MITACS Research Training Award. Wilfrid Laurier University is located on the shared traditional territory of the Neutral, Anishinaabe and Haudenosaunee peoples. This land is part of the Dish with One Spoon Treaty between the Haudenosaunee and Anishinaabe peoples.

## Competing Interests

The authors have no relevant financial or non-financial interests to disclose.

## Authors’ contributions

Karen Liu: Formal analysis, Investigation, Methodology, Writing - review & editing Vanessa Shivnauth: Methodology, Writing - review & editing Christian Danve M. Castroverde: Conceptualization, Funding acquisition, Supervision, Formal analysis, Validation, Writing - original draft, review & editing

## Notes

### Competing Interest Statement

The authors have declared no competing interest.

